# Intrinsic growth heterogeneity of mouse leukemia cells underlies differential susceptibility to a growth-inhibiting anticancer drug

**DOI:** 10.1101/2020.07.10.196881

**Authors:** Akihisa Seita, Hidenori Nakaoka, Reiko Okura, Yuichi Wakamoto

## Abstract

Cancer cell populations consist of phenotypically heterogeneous cells. Growing evidence suggests that pre-existing phenotypic differences among cancer cells correlate with differential susceptibility to anticancer drugs and eventually lead to a relapse. Such phenotypic differences can arise not only externally driven by the environmental heterogeneity around individual cells but also internally by the intrinsic fluctuation of cells. However, the quantitative characteristics of intrinsic phenotypic heterogeneity emerging even under constant environments and their relevance to drug susceptibility remain elusive. Here we employed a microfluidic device, mammalian mother machine, for studying the intrinsic heterogeneity of growth dynamics of mouse lymphocytic leukemia cells (L1210) across tens of generations. The generation time of this cancer cell line had a distribution with a long tail and a heritability across generations. We determined that a minority of cell lineages exist in a slow-cycling state for multiple generations. These slow-cycling cell lineages had a higher chance of survival than the fast-cycling lineages under continuous exposure to the anticancer drug Mitomycin C. This result suggests that heritable heterogeneity in cancer cells’ growth in a population influences their susceptibility to anticancer drugs.

## Introduction

Phenotypic heterogeneity in a cancer-cell population is often linked to differential drug susceptibility and can result in incomplete tumor remission. Although various mutational causes of heterogeneous susceptibility to drugs have been documented [1–3], growing evidence indicates that phenotypic heterogeneity arising due to non-genetic causes also affects survival of individual cancer cells upon drug exposure [4–12]. For example, Sharma, *et al.* showed that a human adenocarcinoma cell line (PC9) produces drug-tolerant persisters (DTPs) that remain viable against lethal doses of anticancer drugs [4]. Shaffer, *et al.* also demonstrated that the survival of a few melanoma cells in a drug-exposed population is correlated with the temporal over-expression of resistance-marker genes [5]. Although the underlying molecular mechanisms may differ depending on the drugs and cancer-cell types, phenotypic cell-to-cell variation might be a general phenomenon underlying the appearance of survivors from an isogenic population of cancer cells exposed to lethal stress [9].

The drug tolerance of individual cell lineages could be correlated with physiological states that exist prior to the drug exposure. In fact, it has been shown that cancer-cell populations whose growth is suppressed, due to their entry into the stationary phase [8], or due to prior exposure to growth-inhibiting chemicals [13, 14], produce significantly higher frequencies of survivors than do actively growing populations. It is also demonstrated that a rare, long-term dormant subpopulation exists in acute lymphoblastic leukemia cells and resists to anticancer treatment [15].

Despite the accumulating evidence on the relationship between the heterogeneity of growth phenotype and drug susceptibility, it remains unexplored what causes such heterogeneity in a cancer cell population. One plausible cause is inhomogeneous environments around individual cells that are omnipresent *in vivo*. For example, the spatial organization of tumor tissue inevitably creates environmental heterogeneity around individual cells [16, 17]. Another but not mutually exclusive possibility is intrinsic fluctuations of cellular states arising from the factors such as stochasticity in gene expression and metabolism [18–20]. Therefore, it is important to examine the extent of phenotypic heterogeneity arising intrinsically even under constant environments and whether it is sufficient to alter individual cells’ drug susceptibility. However, quantifying the properties of intrinsic phenotypic heterogeneity in isolation from externally-driven heterogeneity has been a challenge in cancer cell research.

In this report, we measured the fluctuations of generation time (interdivision time) of individual mouse lymphocytic leukemia cells (L1210) over tens of generations in constant environments using a microfluidic device. We found that a stably maintained culture of L1210 cell line harbors rare, slow-cycling cell lineages. The generation time was positively correlated between mother and daughter cells, which thus revealed the heritability of growth phenotypes. We tested the susceptibility of the slow-cycling cell lineages to an anticancer drug, Mitomycin C (MMC), and found that they tended to survive longer than fast-cycling cell lineages. Therefore, our results highlight that intrinsic heterogeneity in cancer cells’ growth could generate a spectrum of drug sensitivity in a population and may reduce the efficiency of anticancer treatment.

## Results

### L1210 cells can grow and divide stably in the microfluidic device

To monitor single L1210 cells for multiple generations, we fabricated a mammalian-optimized version of mother machine microfluidic device [8, 21–23](Fig 1A-C). The mother machine was originally developed to analyze single bacterial cells [21] and then adapted for studying eukaryotic cells [8, 22, 23]. In these devices, individual cells are placed at the closed end of the growth channels. Growth and division of the cells push them towards the open end, eventually excluding them from the growth channel (S1 Video). An almost identical device was used previously for studying L1210 cells in a CO_2_-independent medium (L-15) [8]. In our present work, we placed the device inside an on-stage CO_2_ chamber and flowed standard CO_2_-dependent medium (RPMI-1640) through the device. We confirmed that the pH of the medium in the device was maintained at 7.5 and that it stayed robust to changes in the flow rate (S1 Fig; [24]).

**Fig 1.**
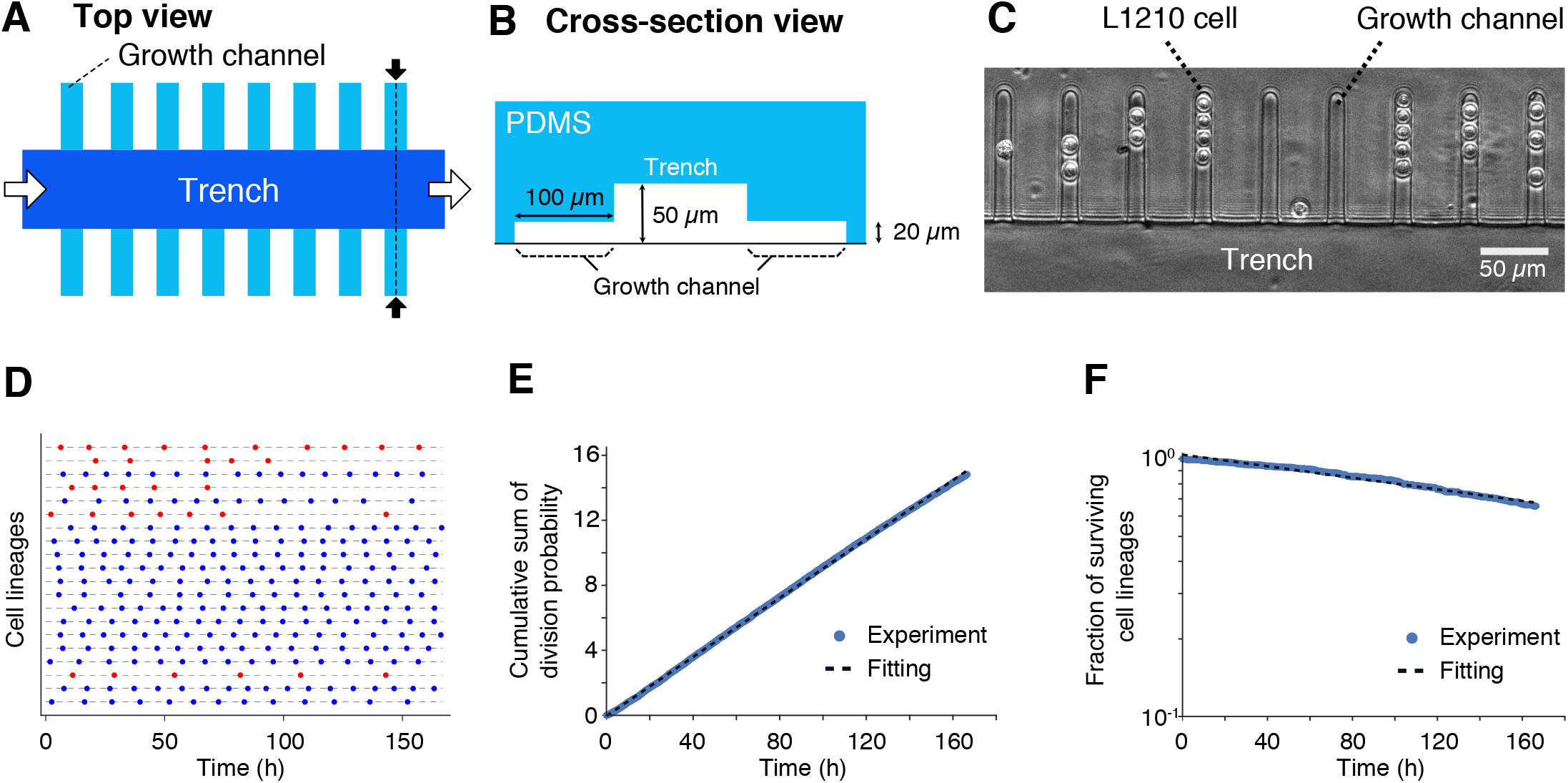
Balanced growth of L1210 cells in the mother machine microfluidic device. (A and B) Schematic representation of the device. (A) The top view of the device. The culture medium flows through the trench (white arrows). Cells trapped in the growth channels are observed simultaneously by time-lapse imaging. (B) A cross-section of the device at the plane corresponding to the broken line indicated by the black arrows in (A). The height and the width of a growth channel are both 20 *μ*m, which corresponds to the size of L1210 cells. (C) A micrograph of L1210 cells in the device. (D) Tracing individual L1210 cell lineages. Twenty representative single-cell lineages of L1210 cells cultured in a constant environment are shown. Each horizontal line represents a single-cell lineage, and the points represent the time points at which cell division occurred. The blue cell lineages are fast-cycling cell lineages, which divided 12 times or more during the seven-day culture; the red cell lineages are slow-cycling cell lineages, which divided 11 times or less during the same period. Co-existence of the heterogeneous growth phenotype is further discussed in the main text. (E) Constant division rate. At each time point *t*, we estimated an instantaneous cell division probability as 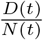, where *N*(*t*) is the number of surviving cells, and *D*(*t*) is the number of cells that underwent cell division within the time-lapse interval (*t*, *t* + Δ*t*]. Δ*t* is 10 min throughout this study. The vertical axis is the cumulative sum of instantaneous division probability, i.e., 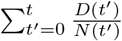. Blue points represent the experimental data. The black broken-line is the line of linear fit; the slope is (9.033 ± 0.003) × 10^−2^ h^−1^. The cumulative sum of division probability increases linearly with *t*, which indicates that the division rate was constant throughout the observation period. (F) Constant death rate. The fraction of surviving cell lineages was plotted over time with the vertical axis on the log scale. Blue points are the experimental data. Black broken-line is the line of linear fit; the slope is −(2.239 ± 0.009) 10^−3^ h^−1^. The surviving fraction decreased in an exponential manner, which indicated a constant death rate throughout the observation period.

We observed hundreds of individual cell lineages simultaneously for one week by time-lapse microscopy and recorded every cell division event (Fig 1D and S1 Video). To check the stability of the growth of the cells in the device, we measured their division rate and found it to be almost constant (9.0 × 10^−2^ h^−1^) throughout observation (Fig 1E). One of the advantages of using the mother machine is that we can analyze death kinetics of a population by simply counting the number of surviving lineages at each time point and plotting a survival curve. Indeed, occasional cell deaths were observed (S1 Video), and the death rate was found to be constant (2.2 × 10^−3^ h^−1^, corresponding to roughly 2% chance of death per generation) (Fig 1F). Cell death under favorable growth conditions with low frequency has also been reported in bacteria and fission yeast [21–23], which presumably reflects an accidental loss of cellular homeostasis or stochastic triggering of signal transduction pathways leading to cell death. The constant division and death rates confirmed that the cells experienced conditions of balanced growth in the microfluidic device.

### Distribution and correlation statistics of generation time

To further characterize the cellular growth phenotype, we plotted the distribution of generation times (Fig 2) and found it to have a mean at 10.40 ± 0.04 h (*n* = 6033). This value is close to the doubling time of L1210 cell populations in batch cultures (11±1 h, *n* = 3), which confirms that the culture conditions in the device were comparable to those experienced in the batch cultures. We observed that the distribution of the generation time of L1210 cells was heavily skewed with a long right tail, which indicates that the L1210 cell population harbors cells that remain undivided for extended periods (Fig 2A and B).

**Fig 2.**
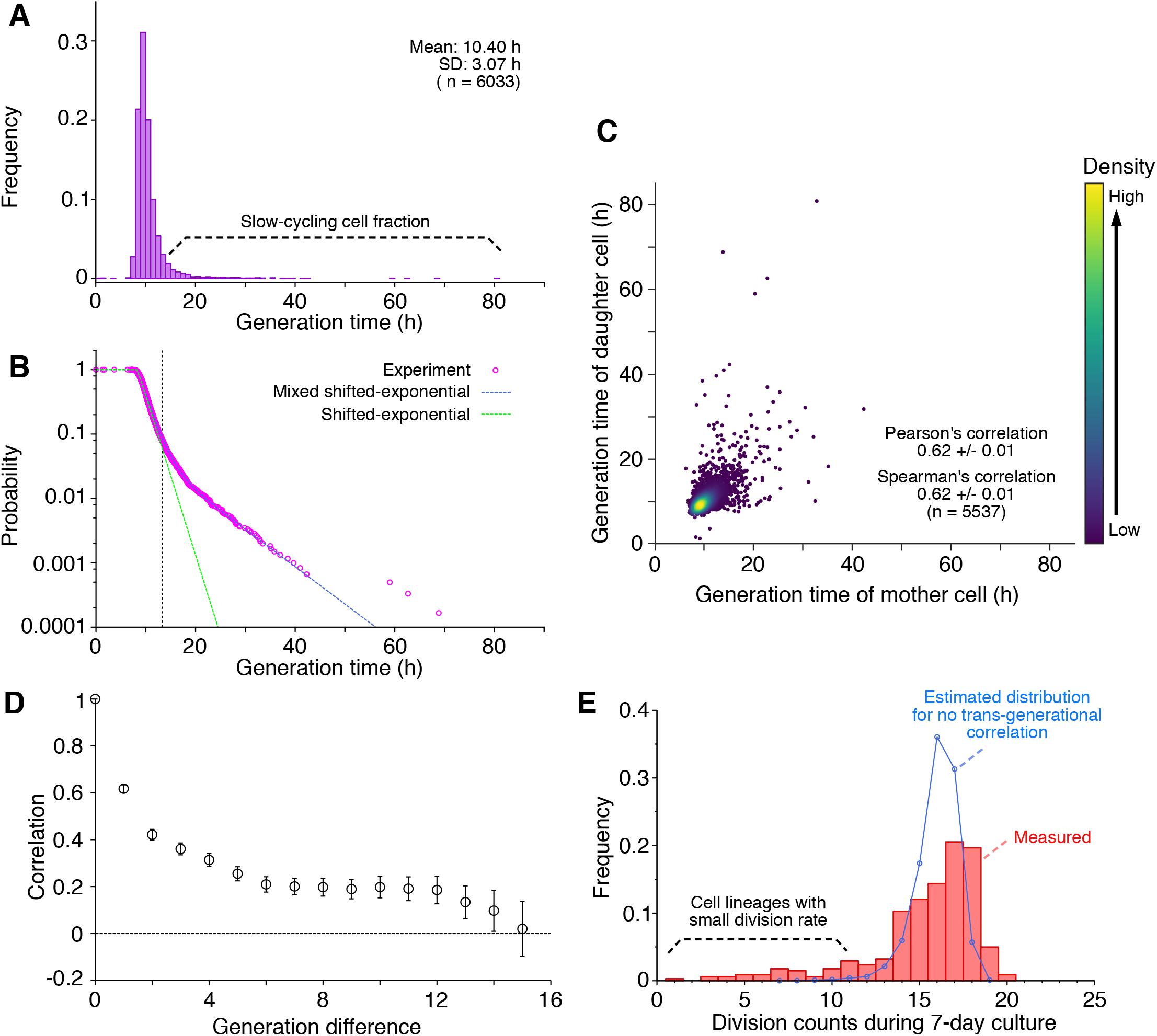
Growth statistics of L1210 cells in a steady environment. (A) Generation time distribution. The mean was 10.40 h, and the standard deviation was 3.07 h (*n* = 6, 033). (B) Survival function of generation time. Magenta points are the experimental data; the blue broken-line represents the curve fitted to the experimental data by a mixed shifted-exponential distribution (Eq 2); the green broken-line represents a curve of a single shifted-exponential distribution. The vertical line at *τ* = 13.31 h indicates the threshold of generation time above which the survival function becomes smaller than 0.06. (C) Correlation between the generation times of mother and daughter cells. (D) Autocorrelation of generation time. Error bars represent standard errors. (E) Division count distribution. The number of cell divisions in each lineage that stayed alive during the seven-day culture (*n* = 341) was counted. The red columns represent the distribution of division counts; the blue curve represents the estimated distribution of division counts, assuming that there is no trans-generational correlation between generation times. The latter was calculated by randomly sampling generation times from the experimental data.

Generation-time distributions of mammalian cells are often fitted to shifted-exponential distributions, whose tails decay with constant rates [25, 26]. However, the observed distribution of the generation times of L1210 cells cannot be fitted to a single shifted-exponential distribution due to an inflection in the decay rate at the generation time of *τ* = 14 h (Fig 2B). Instead, we noticed that the distribution observed for L1210 cells was well-explained by assuming the existence of two subpopulations whose respective shifted-exponential distributions are characterized by distinct decay constants λ_1_ and λ_2_, i.e., the generation time distribution follows the equation:

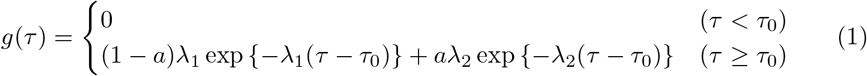

where *τ*_0_ is a fixed no-division period (= minimum cell cycle length), and a (0 < a < 1) is a fraction of cells whose generation time distribution is characterized by the decay constant λ_2_. The probability that a newborn cell remain undivided until age *τ* is given by the survival function (Fig 2):

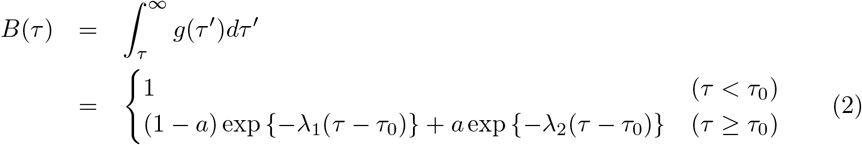

This suggests that the L1210 clonal cell population is composed of multiple (at least two) subpopulations, each of which follows distinct cell division statistics.

We also calculated the correlation between the generation times of mother and daughter cells and found a relatively strong positive correlation (Pearson’s correlation coefficient, *r* = 0.62 ± 0.01; Spearman’s correlation coefficient, *ρ* = 0.62 ± 0.01, Fig 2C). The positive correlation between the generation times of mother and daughter cells observed here is one of the strongest reported to date for eukaryotic and prokaryotic cells [27–33]. The correlation monotonically decayed across generations, but we still found a positive correlation of *r* = 0.20 ± 0.02 even after ten generations (Fig 2D). This result hints at the existence of long-term memory of growth states in individual cells.

We predicted that the existence of a slow-cycling cell subpopulation and the inheritance of generation times should produce a significant heterogeneity in the division frequency among cell lineages. Indeed, we found that the division counts of cell lineages surviving until the end of the observation period showed a wide distribution with a long left-tail (Fig 2E). Thus, some cell lineages underwent division only rarely despite the presence of favorable growth conditions. To corroborate the significance of the correlation of mother-daughter division statistics within lineages, we generated pseudo-lineage data by random re-sampling from the distribution shown in Fig 2A. We confirmed that the left tail of the corresponding division count distribution was reduced (Fig 2E). Therefore, the heavily right-skewed distribution of generation time and the inheritance of the growth phenotype across generations produce the heterogeneity of division frequency among cellular lineages of L1210 cells.

A previous study on the growth of L1210 cells reported that the transition of generation time across generations is essentially a deterministic process characterized by a small set of parameters [30]. To check whether such a deterministic picture also applies to our cases, we performed a correlation-dimension analysis on the generation time data. However, we did not find a convergence of the embedded dimensions of generation time (S2 Fig). This result suggests that the transition of generation time of L1210 cells observed in our experiments is a stochastic rather than a deterministic process.

### Division frequency before anticancer drug exposure is correlated with drug susceptibility

Having characterized intrinsic heterogeneity in growth dynamics within L1210 cell population, we next explored whether these variations can affect susceptibility to an anticancer drug, MMC. To directly test if the division rate of individual cell lineages correlates with their drug susceptibility, we applied MMC containing media in the mother machine and followed the fates of individual cells. We first confirmed that MMC is effective in our mother machine system; upon switching from the standard culture medium to drug-containing ones, population growth rates started to decrease (Fig 3A). As expected, exposure to a higher dose (200 nM) of the drug resulted in a more significant decline in division probability and earlier cessation of the entire population growth than exposure to a lower dose (50 nM) (Fig 3A). We also observed increases in death rates in response to the drug treatment, but with a noticeable delay (approximately 50 h) (Fig 3B). A time-lapse movie clearly showed that cells retain their ability to divide, albeit marked slow-down in cell cycle progression during the delay period (S2 Video). This observation is consistent with the fact that MMC primarily affects DNA replication and does not directly induce cell death. Regardless of the dosage, about 40% of the population survived after a prolonged exposure (100 h) to the drug (Fig 3B), indicating that the killing activity of MMC is saturated at the drug concentrations in our experiments.

**Fig 3.**
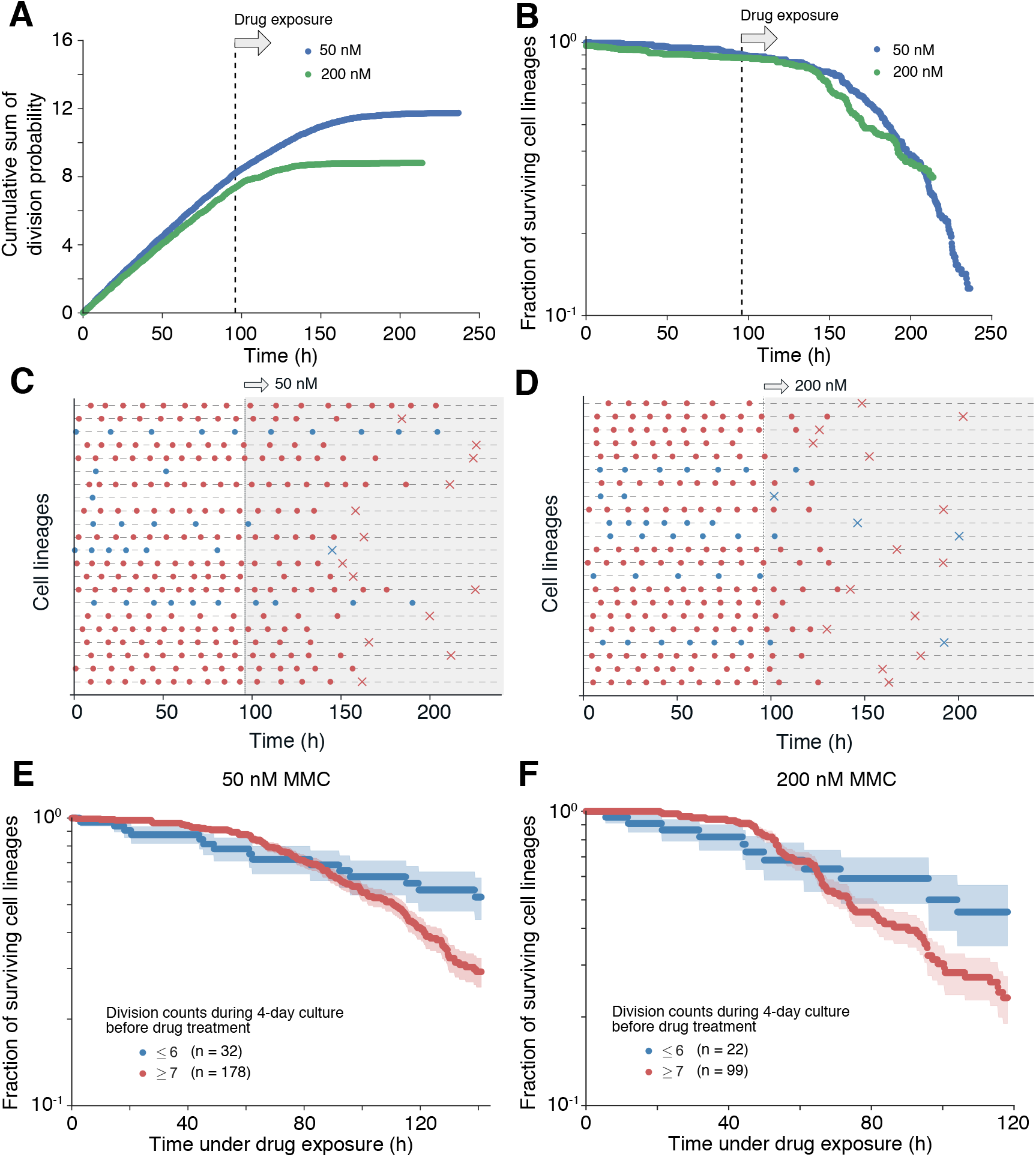
Growth states before drug exposure affect survival. (A) Decrease of division rate upon drug exposure. The cumulative sum of instantaneous division probability of L1210 cells in the microfluidic device was plotted over time. In these experiments, MMC exposure started at 96 h (vertical broken line). Blue points represent the result of exposure to 50 nM MMC and green points represent the result of exposure to 200 nM MMC. In both cases, decreases in the division probability started immediately after the drug exposure (around 100 h). (B) Increase in death rate on drug exposure. The fraction of surviving cell lineages was plotted over time. In both low and high dosage treatment, death rates started to increase around 150 h. (C and D) Division and death events within single-cell lineages. Panel C represents growth under 50 nM MMC, and Panel D represents growth under 200 nM MMC. The gray-shaded time windows represent the periods of drug exposure. Circles represent cell divisions, and crosses cell death. Red lineages represent fast-cycling cell lineages that underwent seven or more divisions during four days of culture before drug exposure; and blue lineages represent slow-cycling cell lineages that underwent six or fewer divisions during the same period. (E and F) Survival curves for fast-cycling (red) and slow-cycling (blue) cell lineages. Shaded regions around the curves represent standard errors. Panel E represents the result of exposure to 50 nM MMC and Panel D represents the result of exposure to 200 nM MMC.

The time points at which the different cell lineages died were largely heterogeneous (Fig 3C and D). Since the division frequencies of the individual cell lineages were also heterogeneous (Fig 1D, 2E, 3C, and 3D), we wanted to examine whether pre-exposure division frequencies correlated with susceptibility to MMC. Therefore, we categorized the cell lineages into two groups: (i) those that underwent seven or more cell divisions during the 96 hours preceding drug exposure and (ii) those that underwent six or fewer cell divisions. The division count cut-off was determined based on the assumption that cells with generation times longer than 14 h were in the slow-cycling state (see Materials and Methods for details).

The cell lineages exhibiting the higher division frequency before drug exposure stayed alive for the first 60 hours of exposure to 50 nM MMC, but the decay of their surviving fraction accelerated thereafter (Fig 3E). On the other hand, we did not observe such acceleration in the death kinetics for the slow-cycling lineages. Consequently, the low and constant death rate allowed the slow-cycling cell lineages to leave a larger fraction of surviving cells (ca. 40%) even after seven days of continuous exposure to MMC (Fig 3E).

Usage of a higher concentration (200 nM) of MMC caused similar decays, but the surviving fraction of the cell lineages exhibiting higher prior division frequencies decayed more rapidly after an initial 40-hour endurance period (Fig 3F). These results suggest that pre-exposure growth states affect the response to MMC treatment and subsequent survival dynamics: Slow growth is detrimental to short-term survival, but advantageous if the drug exposure persists.

## Discussion

Cancer cell populations consist of phenotypically heterogeneous cells. This heterogeneity can influence the effectiveness of anticancer drug treatment and, thus, the chances of relapse. Furthermore, identification of pre-exposure phenotypic characteristics linked to drug resistance might allow us to predict the effectiveness of anticancer drugs even before they are administered. However, understanding what generates the phenotypic heterogeneity among cancer cells is challenging due to the lack of a method to separate the contributions of intrinsic and extrinsic sources of noise.

In this study, we analyzed single L1210 lymphocytic leukemia cells in a strictly-controlled constant environment utilizing microfluidic time-lapse microscopy. This experimental setup enabled us to characterize the intrinsic properties of growth heterogeneity. Statistical analysis of the growth of L1210 cells under favorable culture conditions revealed that the distribution of their generation times was heavily skewed, with a long right-tail. This suggested that the L1210 cell population contained a slow-cycling subpopulation. Besides, the generation times of mother-daughter cell pairs were strongly correlated across generations. The inspection of individual cell lineages for counting division events confirmed that a few lineages undergo division infrequently.

Importantly, the cells present in the slow-cycling state died more gradually upon exposure to MMC (Fig 3), which proves that the intrinsic growth heterogeneity emerging in a constant environment is sufficient to produce variable susceptibility to the drug. Since MMC targets DNA replication, this result might be consistent with the simple view that the lack of DNA replication allows slow-cycling cells to withstand the inhibitory effect by MMC. However, resistance to anticancer drugs may also result from failure to undergo apoptosis or from increased efflux activity [34]. It is also plausible that, in slow-cycling cells, more intracellular resources are allocated to alternative metabolic processes that alleviate the effects of the drug rather than to the processes of growth and replication.

The trans-generational correlation of generation time of L1210 cells was previously measured by Sandler *et al*. using a different experimental setup [30]. They reported that the correlation between mother and daughter cells’ generation times was nearly zero, but mother and granddaughter cells had a positive correlation. Based on this intriguing result, they developed a deterministic model to explain cell-cycle dynamics across generations. Their model predicts the existence of various mother-daughter correlations in a manner highly dependent on the model parameters and the initial conditions. Therefore, the positive mother-daughter correlation observed in our study does not violate their model. Nonetheless, the embedded dimensions should converge irrespective of the parameter values and initial conditions in the correlation dimension analysis. However, our analysis did not find a convergence of the embedded dimensions (S2 Fig). This may suggest that subtle differences in cultivation conditions and cell lines drastically alter the contribution of stochasticity to the determination of generation times of L1210 cells.

The existence of a small subset of slow-cycling or non-growing quiescent cells has been described in various types of cancer cell populations. The contribution of such cells to anticancer drug resistance and the formation of new tumors is of great concern in modern cancer biology [35–38]. Although the detection of slow/non-growing cells often relies on indirect methods such as the label-retaining assay [35, 37], our microfluidic lineage-tracking strategy provided direct evidence of pre-existing slow-cycling L1210 leukemia cells that last longer upon subsequent drug exposure. It is of note that generation times of the slow-cycling leukemia cells characterized in this study were less than 40 h in most cases; thus, they should still be referred to as proliferative rather than quiescent cells. It is an intriguing question whether the slow-cycling phenotype is a prerequisite for the acquisition of stronger drug resistance, possibly mediated by drastic changes in metabolic states or triggering adaptive mutations [39, 40].

## Materials and methods

### Cell line and culture conditions

L1210 mouse lymphocytic leukemia cells were purchased from ATCC (ATCC CCL-219). The cells were grown in 75 cm^2^ culture flasks in an incubator under a 5% CO_2_ atmosphere at 37°C in RPMI-1640 medium (Wako) supplemented with 10% fetal bovine serum (Biosera).

To visualize the cellular nuclei, the cells were transformed with pcDNA3.1-NLS-mVenus (mVenus N-terminally tagged with a nuclear localization signal peptide) by electroporation (super electroporator NEPA 21; NEPA GENE). Stable expression of mVenus in the established cell lines was confirmed by flow cytometry.

### Measurement of population growth rate in batch cultures

L1210’s population doubling time was evaluated with Cell-Counting Kit-8 (CCK-8: Dojin), following the manufacturer’s instructions. Cells in exponential growth were seeded at initial cell density 1.25 × 10^4^ cells/ml in RPMI-1640 medium supplemented with 10% FBS and cultured under 37°C and 5% CO_2_ conditions. 100 *μ*l of cell suspension was transferred to each of 5 wells in a 96-well plate (Corning), and CCK-8 solution (10 *μ*l/well) was added to the wells. After 2 hours incubation under 37°C and 5% CO_2_ conditions, the absorbance (450 nm) of each well was measured with a plate reader (Molecular Devices). Until the stationary phase, the measurement of the absorbance was performed once a day, and the population doubling time was calculated.

### Fabrication of mammalian mother machine microfluidic device

A microfluidic device was fabricated using standard photolithography techniques. First, to make the growth channels, the photoresist SU-8 3025 (MicroChem Corp.) was cast on a silicon wafer (ID 447, *ϕ* = 76.2 mm, University Wafer) using a spin coater (MS-A150, Mikasa) under the following conditions:

1. The spin speed was ramped up to 500 rpm at an acceleration of 100 rpm/sec and then held constant for 5 sec.
2. The spin speed was ramped up to 3,000 rpm from 500 rpm at an acceleration of 100 rpm/sec and then held constant for 30 sec.
3. The spin speed was ramped down to 0 rpm at a deceleration of 100 rpm/sec.

The SU-8-coated silicon wafer was transferred onto a hot plate and baked for 30 min at 95°C (soft bake). After soft bake, patterns of growth channels on a photomask were transferred to the photoresist with a mask aligner (MA-20, Mikasa) using a long-pass filter (LU0350, cut-on wavelength = 350 nm, Asahi Spectra Co.). The wafer was then transferred onto a hot plate and baked for 10 min at 95°C (post-exposure bake). After it had cooled down to room temperature, the unhardened photoresist was dissolved in SU-8 developer (MicroChem Corp.), and the mold was rinsed with isopropanol (Wako) for 30 sec. Next, we fabricated a trench channel directly on the mold of the growth channels. The SU-8 3050 photoresist (MicroChem Corp.) was coated on the mold of the growth channels under the same conditions as employed during the SU-8 3025 coating. The soft bake was carried out at 95°C for an hour. The trench channel was created with a mask aligner such that the trench and the growth channels were orthogonal to each other. The post-exposure bake, development, and rinsing were carried out as described above.

Following the manufacturer’s instructions, we mixed the PDMS (polydimethylsiloxane) elastomer and curing agents (Sylgard184; Dow Corning Toray, Corp., Ltd.) at a ratio of 10:1. The mixture was poured onto the fabricated mold, degassed in a vacuum desiccator, and then baked at 65°C overnight in an oven. Then the cured PDMS was peeled off carefully from the wafer, and then washed with isopropanol for at least 10 min in an ultrasonic cleaner. The device was dried in an oven at 65°C and bonded irreversibly to a clean coverslip (24 mm × 60 mm, thickness 0.12–0.17 mm, Matsunami) by oxygen plasma processing (FA-1, Samco).

### Time-lapse imaging

Exponentially growing cells were concentrated to approximately 1 × 10^6^ cells/ml by centrifugation. The suspension of the cells was then introduced into the microfluidic device through the inlet hole. To fill the growth channels with cells through gravitational sedimentation, we tilted the device vertically and incubated it for at least 30 min in the incubator. After the cell loading, the device was placed in a stage-top incubator (under 5% CO_2_, at 37°C; Tokai Hit) on an inverted microscope (Eclipse Ti, Nikon) equipped with a YFP-HQ filter cube (excitation filter 500/20 nm, dichroic mirror 515 nm, barrier filter 535/30 nm; Nikon). The culture medium (RPMI-1640 supplemented with 10% fetal bovine serum) was supplied continuously through the trench channel using a syringe pump at a flow rate of 1.0 ml/h. This pumping rate was optimized to maintain the pH of the medium flowing through the device for stable cell growth (S1 Fig). To observe the cellular response to MMC, we replaced the drug-free medium with the drug-containing medium (RPMI-1640 with 10% fetal bovine serum and either 50 nM or 200 nM of MMC; Wako). During all the experiments, imaging was performed using a 20× dry objective lens (S plan fluor NA 0.45; Nikon) and a digital CCD camera ORCA R2 (Hamamatsu photonics) at 10 min intervals. The exposure times were 10 ms for bright field and 1000 ms for fluorescence imaging, respectively. LED light (Cold White LED Array Light Sources; Thorlabs) was used for fluorescence excitation.

### Data processing and statistical analysis

We tracked the images manually using ImageJ [41] to record cell division and death events. We judged a cell dead when its morphology collapsed or when the fluorescence signal of mVenus was lost (S3 Fig). The processing of the lineage data set and statistical analysis were performed using Python scripts with the Scipy library or C programs. Incomplete generations (the first and the last cells in each single-cell lineage) were excluded from the statistical analyses.

To fit the survival function of generation times (Fig 2B) using Eq 2, we first determined λ_1_ and λ_2_. We estimated λ_1_ and λ_2_ by fitting an exponential function to the 10-12 h window and the 25-42 h window of the survival function, respectively, using the Marquardt-Levenberg fitting algorithm. We then estimated *a* and *τ*_0_ by fitting Eq 2 to the experimental survival function again by using the Marquardt-Levenberg fitting algorithm. The estimated values of these parameters are λ_1_ = 0.571 ± 0.002 h^−1^, λ_2_ = 0.134 ± 0.003 h^−1^, *τ*_0_ = 8.375 ± 0.002 h, and *a* = 0.060 ± 0.001.

The estimated value of a suggests that approximately 6% of the cells in the population are in the slow-cycling state. We designated *τ* = 14.0 h as the cut-off generation time since *B*(*τ*) ≤ 0.06 when *τ* ≥ 14.0 h. In the experiments in Fig. 3, we observed the cells for 96 h in the medium without the drug before changing it to the one containing MMC. Because 96h/14h = 6.86, we assumed the cell lineages that underwent seven or more divisions before the drug exposure as fast-cycling cell lineages and those that underwent six or less divisions as slow-cycling cell lineages.

### Correlation dimension analysis

To determine whether the generation time dynamics were controlled by stochastic or deterministic processes, we employed the Grassberger–Procaccia algorithm (G-P algorithm) [42] on the series of generation times obtained for each cell lineage. As the G-P algorithm is generally applied to a sufficiently long time series, we assumed ergodicity for the series of generation times of each of the surviving lineages, i.e., each surviving lineage is considered as a part of a (hypothetical) long lineage. The generation-time series obtained from *j*-th (1 ≤ *j* ≤ 301) surviving lineage is represented as **T**(*j*) = {*T*_1_, *T*_2_, … *T*_*n*−1_, *T*_*n*_}, where *T*_*k*_ is the *k*-th generation time in the lineage, and n is the number of complete cell cycles observed in the lineage. Several one-dimensional time series **T**(*j*) can be embedded in an m-dimensional space (1 ≤ *m* ≤ *n*) by constructing *m*-dimensional vector **T**^*m*^(*j*) = (*T*_1_, *T*_2_, …, *T*_*m*_), using the first m entries of **T**(*j*). A set of points **T**^*m*^(*j*) represents an attractor of a dynamical system that rules the generation of the time series, and its correlation dimension *D* can be estimated with the correlation integral *C*^*m*^(*r*), which is defined as

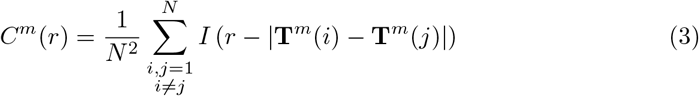

where *N* is the total number of surviving cell lineages, and *I*(·) is the Heaviside function that takes either the value 0 or 1:

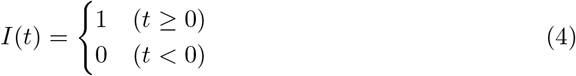

If the generation time series are produced by a stochastic process, then, *C*^*m*^(*r*) ~ *r*^*m*^ for all *m*, whereas *C*^*m*^(*r*) ~ *r*^*D*^ for *m* ≥ *D* if the process is deterministic. For each embedded dimension *m* (1 ≤ *m* ≤ 9), the *C*^*m*^(*r*) of generation-time series data was calculated for varying *r*, and log *C*^*m*^(*r*) was plotted against log *r*. For each plot, local linear fitting on 300 successive data points was carried out by using ranges of five data points (moving linear regression), and the maximum value of the slopes of the fitted lines was reported as the estimated correlation dimension.

## Supporting information

S1 Video. Growth and division of L1210 cells in the mammalian mother machine microfluidic device (S1Video.mp4).

S2 Video. Growth and division of L1210 cells and the response to 200 nM MMC (S2Video.mp4).

**S1 Fig.**
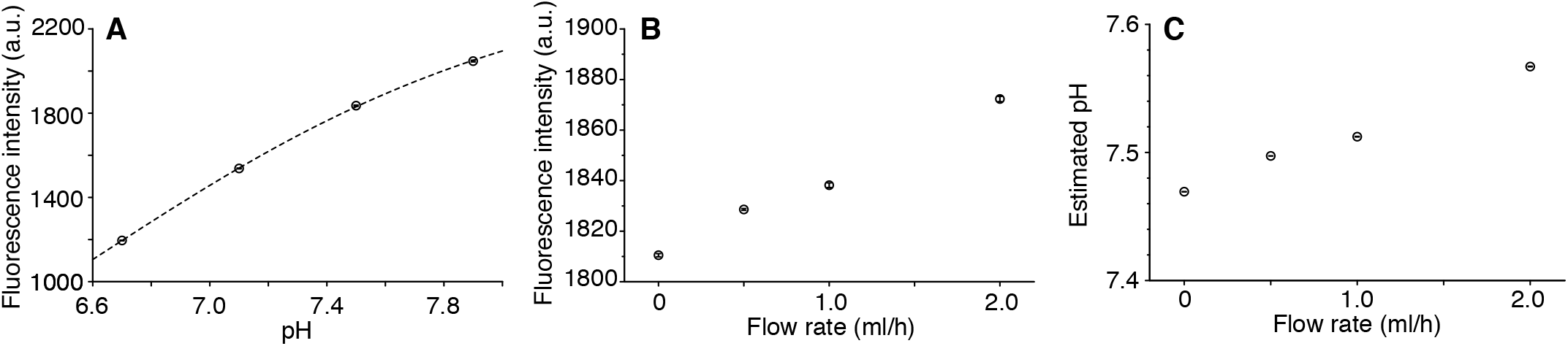
Estimating the pH of the flowing media in the microfluidic device. (A) Fluorescein fluorescence is dependent on pH. 0.1 mM fluorescein solution buffered with 100 mM HEPES (pH 6.7, 7.1, 7.5, or 7.9) was introduced into the device, and fluorescence images in the trench were acquired by microscopy. The points show the measured relationship between the pH of the flowing solution and the fluorescence intensity. The error bars, which are smaller than the points, were the standard deviation of the fluorescent intensity of the images acquired at five times with a 1-min interval at the same position in the trench channel. The broken curve represents the fitting of the data points by a Hill function 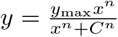 with the parameters *y*_max_ = (2.53 ± 0.05) × 10^3^ a.u., *C* = 6.78 ± 0.03, and *n* = 9.5 ± 0.3. (B) pH of the culture medium in the microfluidic device is robust to fluctuation in the medium flow rate. RPMI-1640 medium containing 0.1 mM fluorescein as a pH reporter was introduced into the device under several conditions of flow rate. The device was placed in the 5% CO_2_ atmosphere on the microscope stage, and fluorescence images were acquired. The plot shows the relationship between the flow rate of RPMI-1640 medium and fluorescence intensity. (C) Relationship between flow rate and pH of RPMI-1640 medium estimated from the results in (A) and (B).

**S2 Fig.**
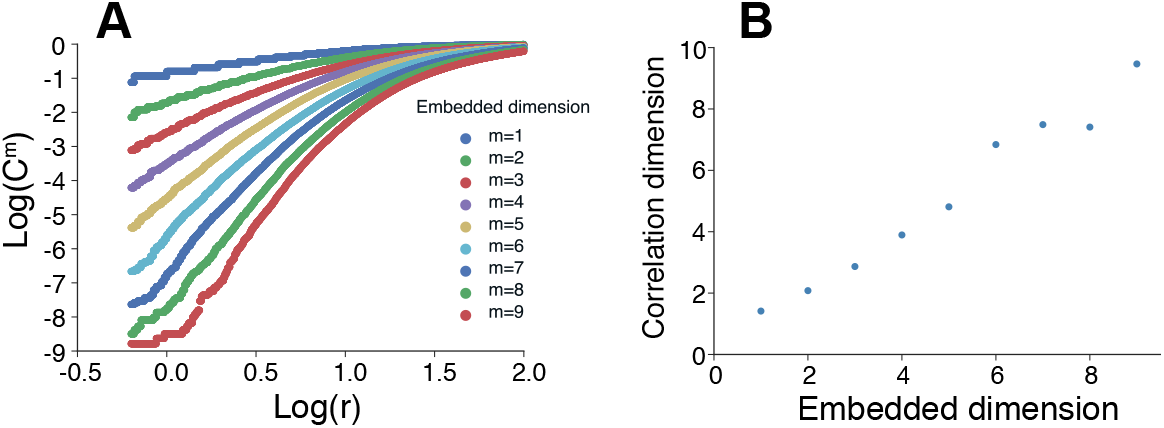
Correlation dimension analysis with the embedded dimension changed from 1 to 9. (A) Correlation integral *C*^*m*^(*r*) plotted against *r* on log-log scale for the experimental data. Correlation integral was calculated based on 301 cell lineages of more than 11 generations. (B) Correlation dimension plotted against the embedded dimension for the experimental data. The correlation dimension with each embedded dimension was estimated as the maximum slope of log *C*^*m*^(*r*) versus log *r* plot in Panel A.

**S3 Fig.**
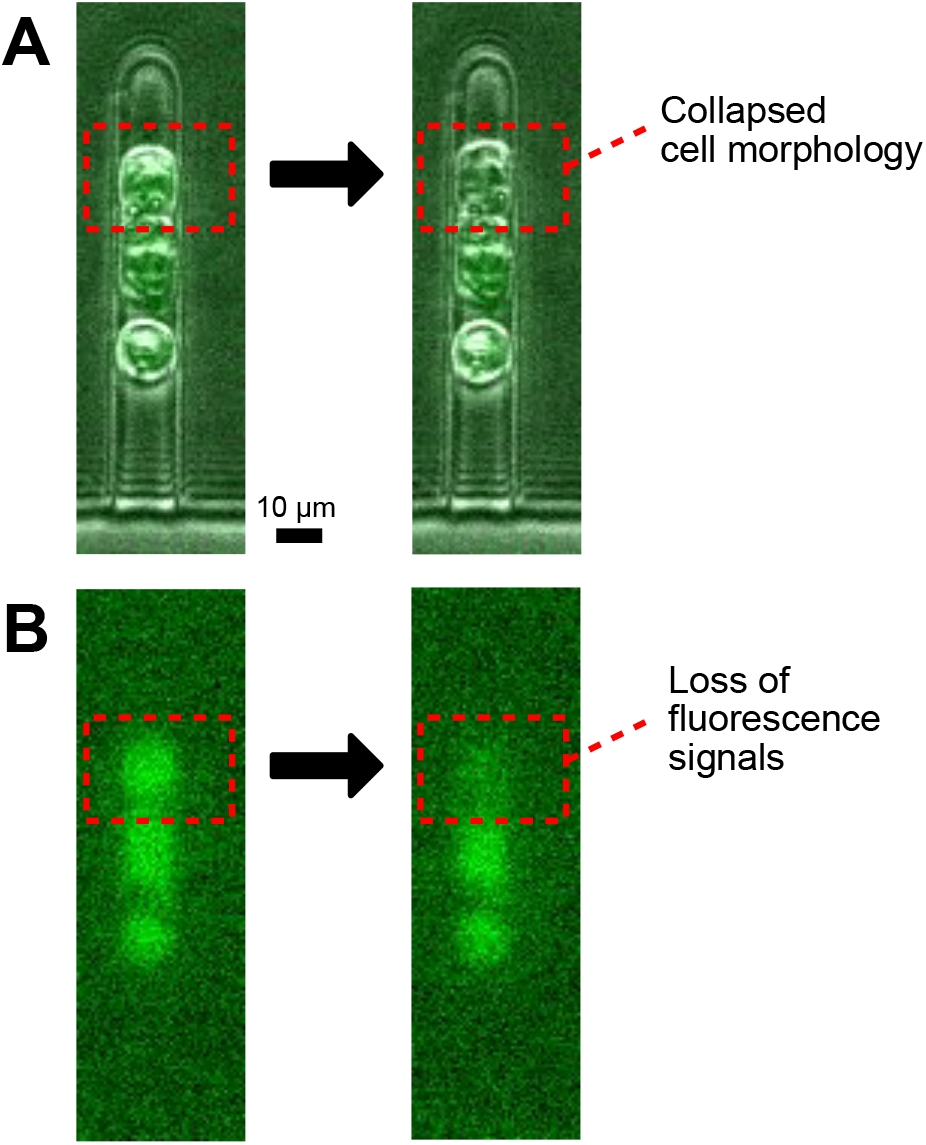
Detection of death events. (A) Microscopic images of a cell exhibiting death in a growth channel. An intact-looking cell in the red rectangular collapsed its cell morphology by the next time point in the time-lapse measurement. Since these collapsed cells stopped moving and never regrew, we judged them dead. (B) The fluorescence images of a cell. The images correspond to those in A. The mVenus signal in the cell dropped significantly by the next time point. Since these cells stopped moving and never regrew, we also regarded the loss of mVenus fluorescence signals as the indication of cell death.

## Acknowledgments

We thank the members of the Wakamoto lab for discussions. This work was supported by JST CREST Grant Number JPMJCR1927 (to Y.W.); and Japan Society for the Promotion of Science KAKENHI (grant number 15KT0075, 15H05746, 17H06389, and 19H03216 to Y.W.).

